# *Krüppel-like factor* gene function in the ctenophore *Mnemiopsis leidyi* assessed by CRISPR/Cas9-mediated genome editing

**DOI:** 10.1101/527002

**Authors:** Jason S Presnell, William E Browne

## Abstract

The *Krüppel-like factor* (*Klf*) gene family encodes for transcription factors that play an important role in the regulation of stem cell proliferation, cell differentiation, and development in bilaterians. While *Klf* genes have been shown to functionally specify various cell types in non-bilaterian animals, their role in early diverging animal lineages has not been assessed. Thus, the ancestral activity of these transcription factors in animal development is not well understood. The ctenophore *Mnemiopsis leidyi* has emerged as an important non-bilaterian model system for understanding early animal evolution. Here we characterize the expression and functional role of *Klf* genes during *M. leidyi* embryogenesis. Zygotic *Klf* gene function was assessed with both CRISPR/Cas9-mediated genome editing and splice-blocking morpholino oligonucleotide knockdown approaches. Abrogation of zygotic *Klf* expression during *M. leidyi* embryogenesis results in abnormal development of several organs including the pharynx, tentacle bulbs, and apical organ. Our data suggest an ancient role for *Klf* genes in regulating endodermal patterning, possibly through regulation of cell proliferation.

**Summary Statement:** Using CRISPR/Cas9 genome editing and morpholino oligonucleotide knockdown, this study shows that tissues derived from the endoderm are dependent upon *Klf5* ortholog expression for proper development and patterning in the ctenophore *Mnemiopsis leidyi*.

## Introduction

Members of the *Krüppel-like factor (Klf)* gene family encode transcription factors with a characteristic DNA binding domain composed of three C-terminal C2H2-zinc fingers (McConnell and Yang, 2010; Presnell et al., 2015). During metazoan diversification, the *Klf* transcription factor gene family expanded via duplication and domain shuffling events (Presnell et al., 2015). *Klf* transcription factors are expressed in a variety of cells and tissues and have roles in many biological processes, including proliferation of stem and progenitor cells, embryonic development, germ layer differentiation, neuronal growth and regeneration, immune system regulation and metabolic regulation (Bialkowska et al., 2017; McConnell and Yang, 2010; Moore et al., 2009; Nagai et al., 2009; Oishi and Manabe, 2018; Pearson et al., 2008; Sweet et al., 2018).

While KLF functional studies have been restricted to bilaterians, *Klf* genes are found in the genomes of all metazoans (Presnell et al., 2015) with a number of homologs expressed in multipotent stem cells (Tarashansky et al., 2021). Within Cnidaria, *Klfs* are expressed in multipotent interstitial stem cells and their various downstream lineages, as well as in ectodermal epithelial stem cells in *Hydra vulgaris* (Hemmrich et al., 2012; Levy et al., 2021; Siebert et al., 2019). In cnidarian single-cell RNA-seq datasets, *Klfs* are expressed in various cell types, including gastrodermis, neuronal and gland cell lineages (Levy et al., 2021; Sebé-Pedrós et al., 2018a). In *Hydractinia symbiolongicarpus, Klf* genes are upregulated in male sexual polyp bodies vs female sexual polyp bodies (DuBuc et al., 2020). Within Porifera, *Klfs* are expressed in the stem-cell like archaeocytes, epithelial pinacocytes, and mesenchymal cells in both *Spongilla lacustris* and *Amphimedon queenslandica* (Musser et al., 2019 preprint; Sebé-Pedrós et al., 2018b). Single-cell RNA-seq data for the Placozoan *Trichoplax adhaerens* revealed a single *Klf* gene expressed in epithelial cells (Sebé-Pedrós et al., 2018b). In ctenophores, three *Klf* genes have been identified in two distantly related species, *Pleurobrachia bachei* and *Mnemiopsis leidyi* (Presnell et al., 2015). The genome of *M. leidyi* contains *MleKlf5a, MleKlf5b*, and *MleKlfX* (Presnell et al., 2015). *MleKlf5a* and *MleKlf5b* are the result of a lineage-specific duplication within the Ctenophora, while *MleKlfX* is highly derived with no clear orthology to any known metazoan *Klf* clade (Presnell et al., 2015). To date, single-cell and tissue-specific RNA-seq studies in *M. leidyi* have not established differential expression signatures for *Klf* genes (Babonis et al., 2018; Sebé-Pedrós et al., 2018b).

*M. leidyi* is a species of the non-bilaterian phylum Ctenophora, one of the earliest-diverging extant metazoan lineages (Dunn et al., 2008; Hejnol et al., 2009; Kapli and Telford, 2020; Li et al., 2021; Shen et al., 2017; Whelan et al., 2017). *M. leidyi* has been used extensively as a model for investigating early metazoan developmental patterning, regeneration, and the evolution of animal traits (Babonis et al., 2018; Bessho-Uehara et al., 2020; Fischer et al., 2014; Martindale and Henry, 1999; Presnell et al., 2016; Reitzel et al., 2016; Salinas-Saavedra and Martindale, 2020; Schnitzler et al., 2014; Yamada et al., 2010). *M. leidyi* embryos undergo a ctenophore-specific early cleavage program with gastrulation taking place ∼3-5 hours post-fertilization (hpf) followed by tissue organization and organogenesis over the next several hours (Fischer et al., 2014; Freeman, 1976; Fig. 1A). Four pairs of ctene rows, one pair in each quadrant, are typically among the first differentiated ectodermal structures to appear (Fischer et al., 2014). Each ctene plate is made up of polster cells bearing fused giant cilia (Tamm, 1973). While initial ctene plate development is established by maternal factors (Fischer et al., 2014), new ctene row expansion begins post-hatching during the juvenile cydippid stage (Tamm, 2012). After the formation of the initial ctene rows, the developing embryo rapidly increases in size. This period of rapid growth is accompanied by pharynx elongation along the aboral/oral axis, the development of tentacle bulbs, and deposition of the first lithocytes onto the balancer cilia of the apical organ (Martindale and Henry, 2015). Lithocytes are mineralized cells that form a statolith housed within the apical organ that functions to control orientation in the water column by coordinating ctene row beating (Jokura and Inaba 2020; Tamm 1973; Tamm 2014).

**Fig 1.**
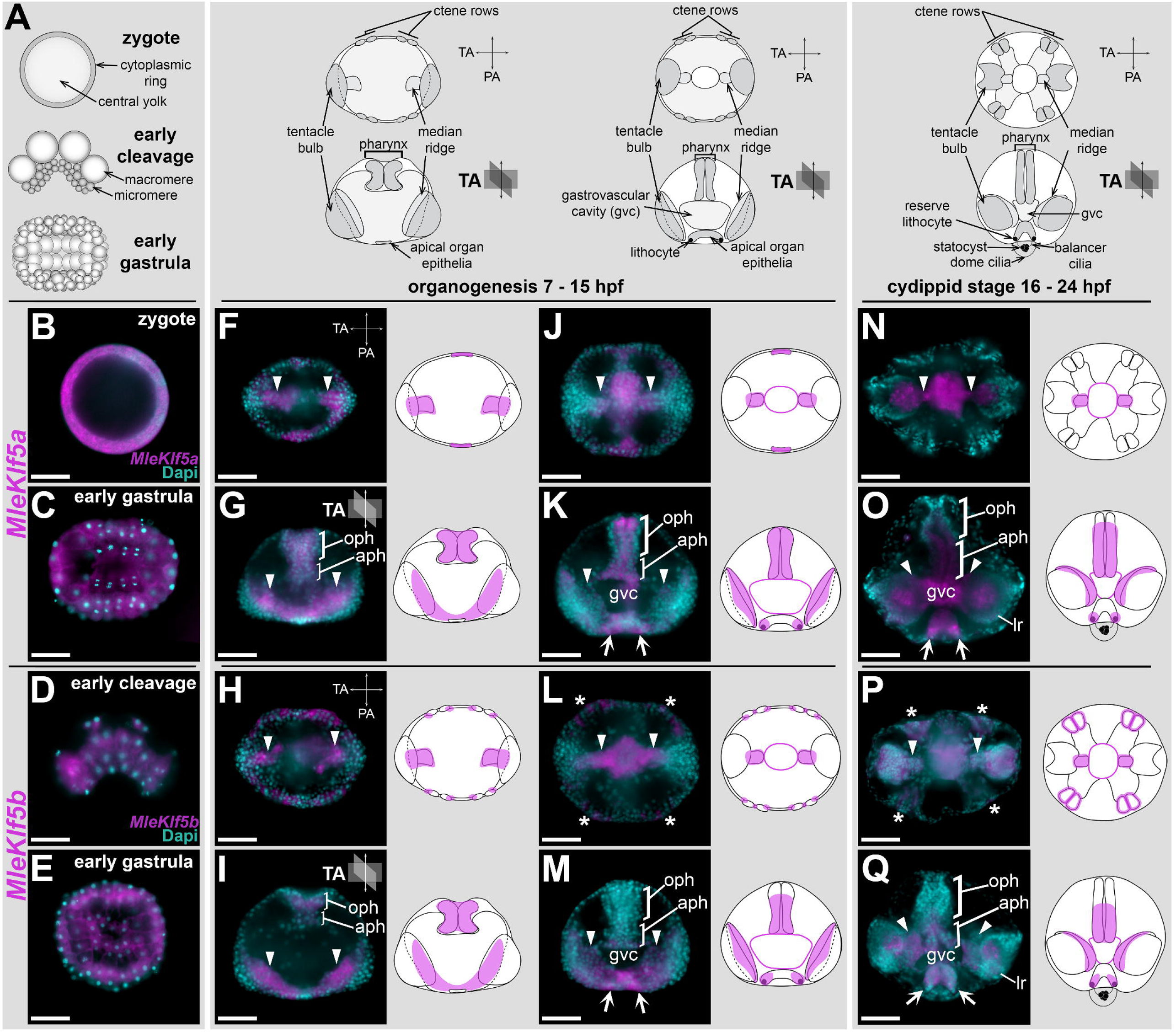
Zygotic *MleKlf5a* and *MleKlf5b* are primarily expressed in endodermally derived tissues during embryogenesis in *Mnemiopsis leidyi*. (**A)** Schematics highlighting major morphological landmarks (e.g., ctene rows, pharynx, tentacle bulbs, apical organ) during *M. leidyi* embryogenesis. Gastrulation typically occurs within 6 hours post-fertilization (hpf), followed by rapid tissue remodeling and organogenesis over the next several hours. By 24 hpf embryos are ready to hatch as cydippid larvae and have fully developed organ systems. For post-gastrulation embryos, the top row is an aboral view and the bottom row is a lateral view with oral up and aboral down. (**B-Q**) Whole-mount *in situ* hybridization for *MleKlf5a* in **B**,**C**,**F**,**G**,**J**,**K**,**N**,**O** and *MleKlf5b* in **D**,**E**,**H**,**I**,**L**,**M**,**P**,**Q** during embryogenesis. Orientation follows schematics from **A**. Aboral views in **C, E, F, H, J, L, N, P**. Lateral views in **D, G, I, K, M, O, Q**. (**B-E**) Maternal transcripts for both *MleKlf5a* and *MleKlf5b* are ubiquitously distributed during early development in zygotes (**B**), early cleavage stages (**D**) and gastrulae (**C**,**E**). One representative image for each gene per stage is shown. (**F-Q**) Zygotic *MleKlf5a* and *MleKlf5b* transcript expression domains with corresponding schematics. (**F-I**) Initially, expression of *MleKlf5a* and *MleKlf5b* zygotic transcripts are localized to the forming tentacular median ridges (arrowheads) and the developing pharynx (oph + aph). (**J-M**) Later in development, *MleKlf5a* and *MleKlf5b* transcript expression are also found in the developing apical organ (arrows) and epithelia of the newly formed gastrovascular cavity (gvc). (**N-Q**) In cydippids, *MleKlf5a* and *MleKlf5b* transcripts are found in the tentacular median ridge (arrowheads) and lateral ridge (lr), on either side of the apical organ floor (arrows), localized towards the aboral end of the pharynx (aph), and throughout the gastrovascular cavity epithelium (gvc). (**L, P**) *MleKlf5b* transcripts are also expressed in an additional domain around the ctene rows (asterisks). See also Fig. S1A-B. Scale bars: 50 µm. aph, aboral end of the pharynx; gvc, gastrovascular cavity; lr, lateral ridge; oph, oral end of the pharynx; PA, pharyngeal axis; TA, tentacular axis.

Flanking the apical organ along the tentacular axis, a pair of ectodermal invaginations and internal endodermal cells form the developing tentacle bulb organs and their cognate tentacular lateral and median ridges, respectively (Martindale and Henry, 1997b; Martindale and Henry, 1999). Embryonic and adult tentacle bulb organs contain populations of highly proliferative cells in the tentacular lateral ridge and median ridge tissues that give rise to differentiated colloblast and tentacle muscle cells, respectively (Alié et al., 2011; Babonis et al., 2018; Jager et al., 2008; Schnitzler et al., 2014). Genes associated with germline development and stemness, including *Piwi, Vasa, Nanos*, and *Sox* homologs are highly expressed in both the lateral and median ridges of the tentacle bulb, as well as in proliferative cell populations in the developing apical organ and ctene rows, supporting the presence of progenitor cells with stem-cell like properties in these tissues. At ∼18-20 hpf, the fully developed *M. leidyi* cydippid hatches and maintains a feeding, pelagic lifestyle before transitioning to the adult lobate body plan ∼20 days post hatching (Martindale and Henry, 2015). In adult animals, these progenitor cells play a role in the continuous replacement of lost cells (Alié et al., 2011; Jager et al., 2008; Reitzel et al., 2016; Schnitzler et al., 2014).

Previous investigations into gene function in *M. leidyi* have utilized morpholino oligonucleotide mediated knockdown or mRNA overexpression methods (Jokura et al., 2019; Salinas-Saavedra and Martindale, 2020; Yamada et al., 2010). Here we report the first use of CRISPR/Cas9 in *M. leidyi* for mutagenesis. We utilized CRISPR/Cas9 to disrupt zygotic function of two *Klf* genes, *MleKlf5a* and *MleKlf5b*. We show that disruption of *Klf* gene expression is associated with the abnormal development of various organs during *M. leidyi* embryogenesis due to the loss of specific endodermally derived cell types. Our data provides additional insight into the evolution of *Klf* gene family function both within the metazoan stem lineage and the early diverging ctenophore lineage. Our use of CRISPR/Cas9 to disrupt *Klf* gene expression and the subsequent characterization of the loss of *Klf* expression on development and tissue patterning in *M. leidyi* provide a foundation for future mutagenesis studies in ctenophores.

## Results

### *MleKlf5a, MleKlf5b*, and *MleKlfX* expression during embryonic development

*MleKlf5a* and *MleKlf5b* transcripts are maternally loaded in *M. leidyi* (Davidson et al., 2017) similar to maternal loading of a number of *Klf* genes in other metazoans (Blakeley et al., 2015; De Graeve et al., 2003; Weber et al., 2014). *MleKlf5a* and *MleKlf5b* transcripts were detected in all embryonic cells through gastrulation (Fig. 1B-E). Post-gastrulation, transcripts for both *MleKlf5a* and *MleKlf5b* became spatially restricted to cell populations associated with the developing pharynx, gastrovascular system, tentacle bulb median ridges, and within the developing apical organ (Fig. 1F-Q).

Within the developing pharynx, *MleKlf5a* and *MleKlf5b* expression were initially widespread (Fig. 1G,I). As the pharynx elongated, *MleKlf5a* and *MleKlf5b* expression became restricted to the interior-most cell layers of the medial and aboral pharyngeal regions (Fig. 1K,M,O,Q). The aboral-most region of the pharynx includes cells that form the junction with the central gastrovascular cavity, or infundibulum. *MleKlf5a* and *MleKlf5b* expression was found throughout the endodermal epithelial lining of the presumptive gastrodermis (Fig. 1J-Q). During the initial development of the aboral apical organ, *MleKlf5a* and *MleKlf5b* expression was detected in the apical organ floor epithelia. As the apical organ developed, *MleKlf5a* and *MleKlf5b* expression became progressively restricted to cells located along the tentacular axis that are positionally correlated with sites of lithocyte formation (Tamm, 2014; Fig. 1K,O,M,Q). Within the developing tentacle bulbs, both *MleKlf5a* and *MleKlf5b* were expressed in the tentacular median ridge (Fig. 1F-Q). An additional unique *MleKlf5b* expression domain was detected in a narrow band of epidermal cells surrounding newly formed ctene row polster cells (Fig. 1H,L,P, Fig. S1A,B).

In contrast to both *MleKlf5a* and *MleKlf5b, MleKlfX* expression was restricted to late embryogenesis, first appearing ∼16 hpf. Expression of *MleKlfX* transcripts were localized to a small number of cells within the apical organ (Fig. S1C,D). One group of *MleKlfX* expressing cells was located deep within the central epithelial floor of the developing apical organ. These cells were located along the tentacular axis and pharyngeal axis forming a cross-shaped pattern (Fig. S1C). A second shallower group of *MleKlfX* expressing cells was located within each quadrant just medial of the ciliated grooves in the developing apical organ (Fig. S1D). These *MleKlfX* expressing cells correspond positionally with the apical organ lamellate bodies, which may represent putative photoreceptor cells (Horridge, 1964a; Jokura and Inaba, 2020; Schnitzler et al., 2012), suggesting that *MleKlfX* expression may be associated with light sensing neuronal cell types in the apical organ.

### CRISPR/Cas9 and splice-blocking morpholino experimental design

To characterize zygotic *Klf* gene function in *M. leidyi*, we used both CRISPR/Cas9 mutagenesis and splice-blocking morpholinos (sbMOs) to independently knockdown *Klf* gene expression during embryonic development (Fig. 2). We focus on *MleKlf5a* and *MleKlf5b* knockdown experiments, as initial *MleKlfX* gene knockdown experiments failed to produce obvious morphological phenotypes. Importantly, it has been previously shown that co-expressed KLFs bind to shared downstream regulatory targets resulting in complex functional outcomes. For example, KLF2, KLF4, and KLF5 have redundant roles in the downstream regulation of *Nanog* (Jiang et al., 2008), KLF2 and KLF4 play redundant roles in regulating attachment between tendon and bone tissue (Kult et al., 2021), whereas competition between KLF1 and KLF3 for binding sites can result in disparate functional outcomes (Ilsley et al., 2017). For this reason, we sought to maximize the efficiency of generating an observable phenotype by performing simultaneous *MleKlf5a* and *MleKlf5b* knockdown with both sbMO and CRISPR/Cas9 genome editing experiments.

**Fig 2.**
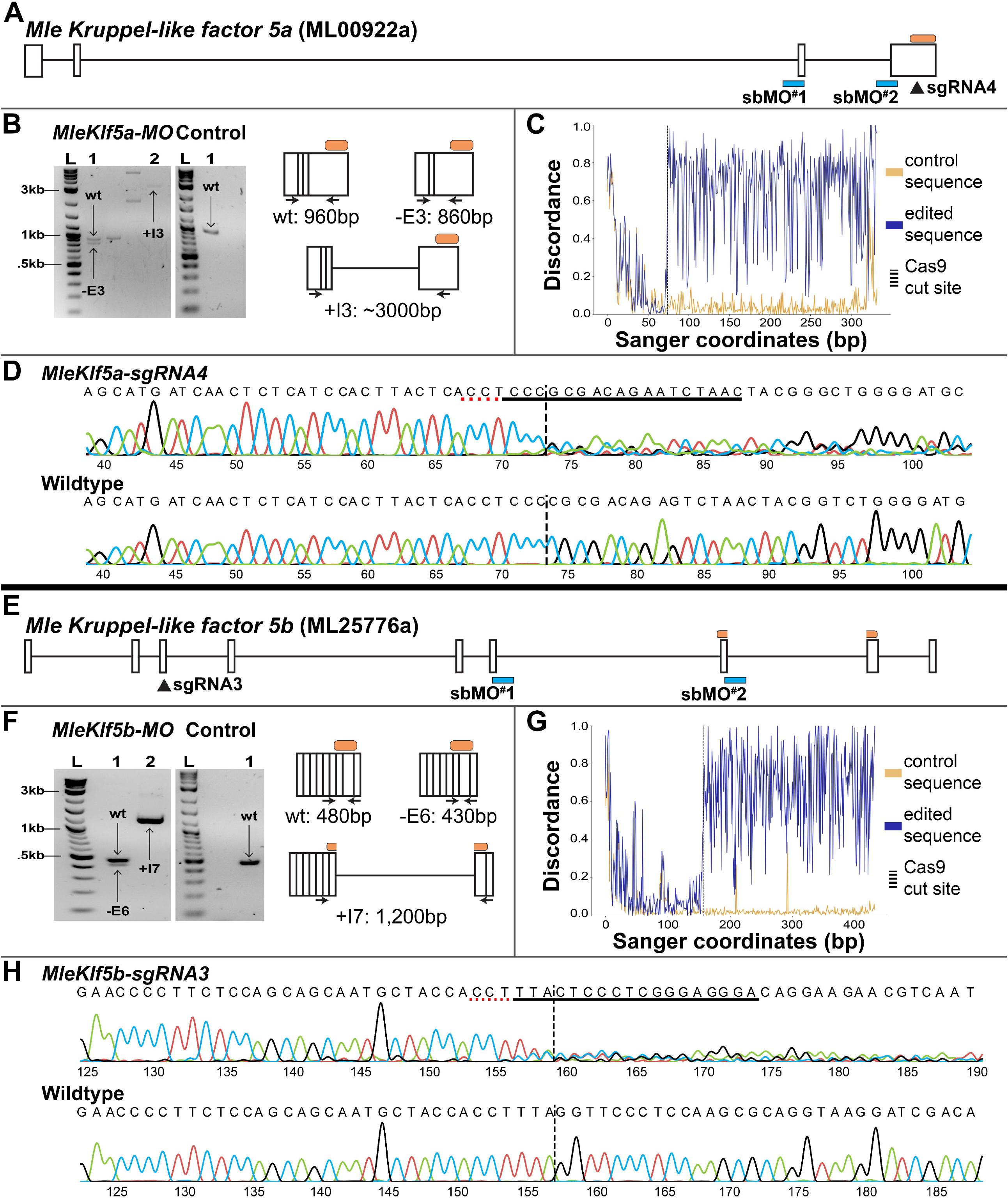
Validation of independent methods used to abrogate *MleKlf5a* and *MleKlf5b* gene function. *MleKlf5a* (**A**) and *MleKlf5b* (**E**) exon-intron schematics show the location of splice-blocking morpholino oligonucleotide (sbMO) targets (blue boxes) and single-guide RNA (sgRNA) targets (black triangles) used in this study. The orange bars indicate the location of the DNA binding domain. (**B, F**) Electrophoretic gels of PCR products obtained using different sets of *MleKlf5a* and *MleKlf5b* sbMO RT-PCR primers on cDNA obtained from a single individual KLF-MO embryo exemplar (left) and control embryo exemplar (right). Schematics to the right of the gel images highlight examples of wildtype (wt) amplicon, exon-skipping (-E) and/or intron retention (+I) amplicons captured with primers for each gene (Table 1). (**B**) *MleKlf5a-MO* gel: 2-log DNA ladder (L) used for band size reference, unlabeled lanes are not relevant to this study. Lane 1 shows both a 960 bp wt and a 860 bp third exon-skipped (-E3) *MleKlf5a* amplicon. Lane 2 shows a ∼3 kb third intron retention (+I3) *MleKlf5a* amplicon. Control gel: 2-log DNA ladder (L) used for band size reference. Lane 1 shows a single 960 bp wt *MleKlf5a* amplicon. **(F)** *MleKlf5b-MO* gel: 2-log DNA ladder (L) used for band size reference. Lane 1 shows both a 480 bp wt and a 430 bp sixth exon-skipped (-E6) *MleKlf5b* amplicon. Lane 2 shows a ∼1.2 kb seventh intron-retained (+I7) *MleKlf5b* amplicon. Control gel: 2-log DNA ladder (L) used for band size reference. Lane 1 shows a single 480 bp wt *MleKlf5b* amplicon. Wildtype (wt) and mis-spliced transcripts due to -E and/or +I were present in KLF-MO embryos (*n* = 21 KLF-MO embryos). (**C-G)** Discordance plots produced using ICE software show elevated sequence discordance downstream of predicted Cas9 cut sites relative to control genomic sequence. (**D, H**) Corresponding Sanger sequence traces from genomic DNA extracted from a single individual exemplar KLF-Cas9 embryo show signal degradation downstream of the Cas9 cut site as compared to a single individual exemplar wildtype embryo (*n* = 17 KLF-Cas9 embryos). Sanger sequencing signal degradation is caused by the introduction of indels in KLF-Cas9 embryos. The sgRNA target sites are underlined, the position of the predicted Cas9 cut sites are represented by a vertical dashed line.

We injected single-cell embryos with either *MleKlf5a*+*MleKlf5b* sbMOs (KLF-MO embryos) or *MleKlf5a-sgRNA*+*MleKlf5b-sgRNA* (KLF-Cas9 embryos). Microinjected embryos were allowed to develop to ∼20 hpf, stained with vital dyes, live-imaged, and compared to equivalent late-stage wildtype embryos from the same spawns. We used fluorescence based vital dyes to mark and follow asymmetries in subcellular components associated with key morphological structures in live animals. For example, MitoTracker fluorescence in whole embryos preferentially marks ctene row polster cells containing giant mitochondria with atypical cristae (Horridge, 1964b). In contrast, LysoTracker fluorescence in whole embryos preferentially marks cells containing yolk and large acidic vacuoles associated with the developing gastrovascular cavity and endodermal canals. After phenotype documentation, live animals were recovered and individually processed for DNA or RNA to validate either CRISPR/Cas9 or MO activity, respectively.

Efficient microinjection of knockdown and knockout reagents requires the mechanical removal of the outer vitelline membrane surrounding the fertilized egg. To determine if mechanically removing the vitelline membrane had an effect on embryogenesis, we scored the percentage of normal development in embryos that were removed from the vitelline membrane but not injected. There was no significant difference in the percentage of normal development between embryos kept in their vitelline membrane (85%, *n* = 161) and those that had their vitelline membrane removed but not subsequently microinjected (80%, *n* = 217; χ^2^ =1.5272, *p*=0.217; Fig. S2A). Additionally, microinjections with a standard control MO (79% normal, *n* = 49), Cas9 protein alone (86% normal, *n* = 7), or with sgRNAs alone (75% normal, *n* = 4) also had no detectable effect on embryonic development (Fig. S2A).

We validated gene expression knockdown efficiency by selecting a subset of KLF-MO, KLF-Cas9, and wildtype embryos for single-embryo RNA or DNA analyses post-experimental manipulation. For both *MleKlf5a* and *MleKlf5b*, sbMOs produced mRNA splicing errors in KLF-MO embryos via exon skipping and/or intron retention (Fig. 2A,B,E,F). An initial set of 4 single guide RNAs (sgRNAs) were designed for *MleKlf5a* and *MleKlf5b* (Table 1) based on the *M. leidyi* reference genome (Moreland et al., 2014; Moreland et al., 2020; Varshney et al., 2015). For each gene, a single sgRNA, *MleKlf5a-sgRNA4* and *MleKlf5b-sgRNA3* (Fig. 2A,E), proved efficient at mediating Cas9 double-stranded break activity at the target loci (Fig. 2C,D,G,H). Sanger sequencing followed by ICE analysis (Hsiau et al., 2019 preprint) revealed a clear degradation of sequence trace signal at the target loci in KLF-Cas9 embryos as compared to control embryos (Fig. 2C,G), indicating the presence of indels and putative frameshift mutations generated by sgRNA targeted Cas9 exonuclease activity (Fig. S2C). ICE analysis predicted the occurrence of frameshift mutations between ∼20-30% (Fig. S2C).

**Table 1.**
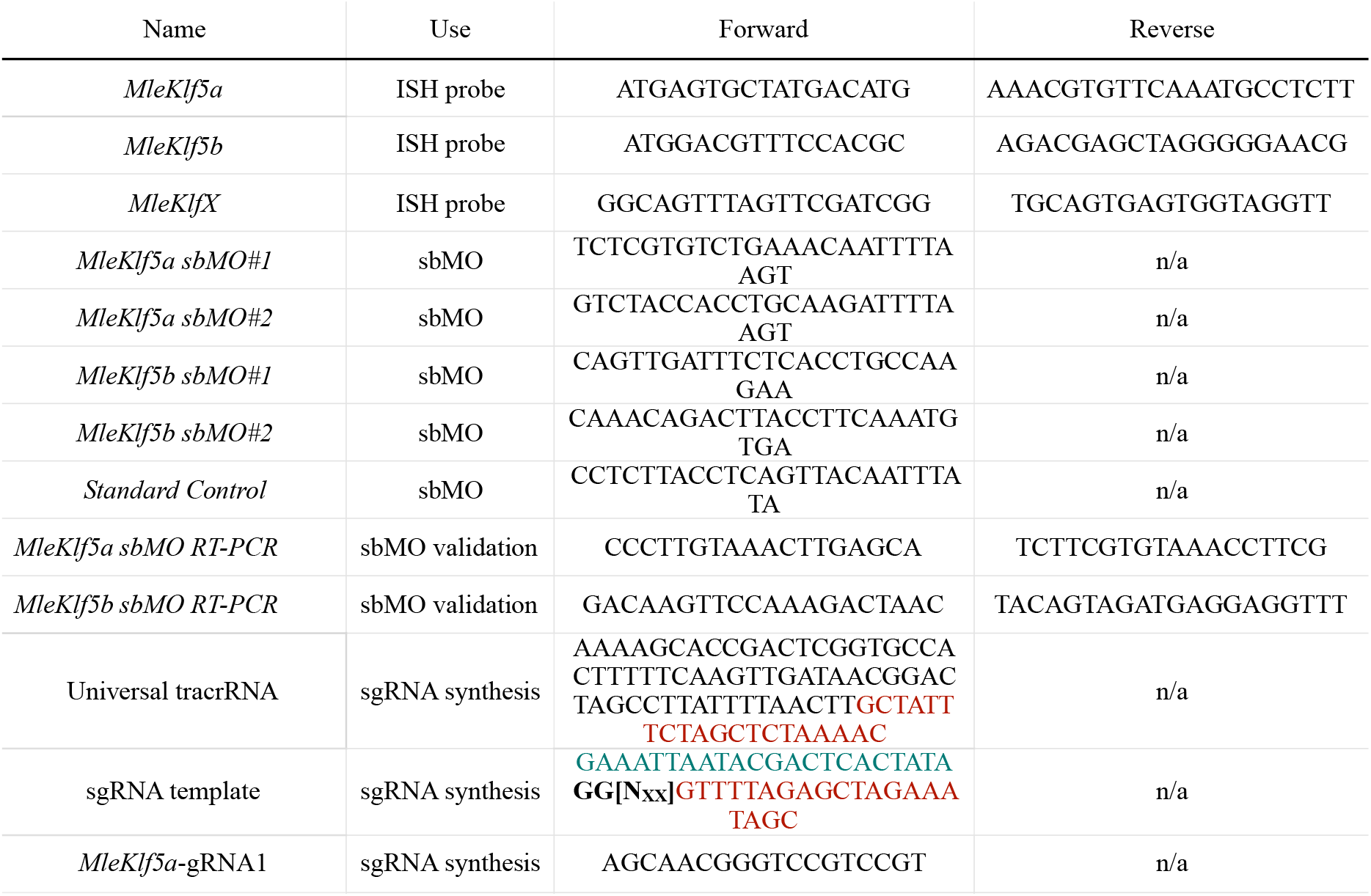
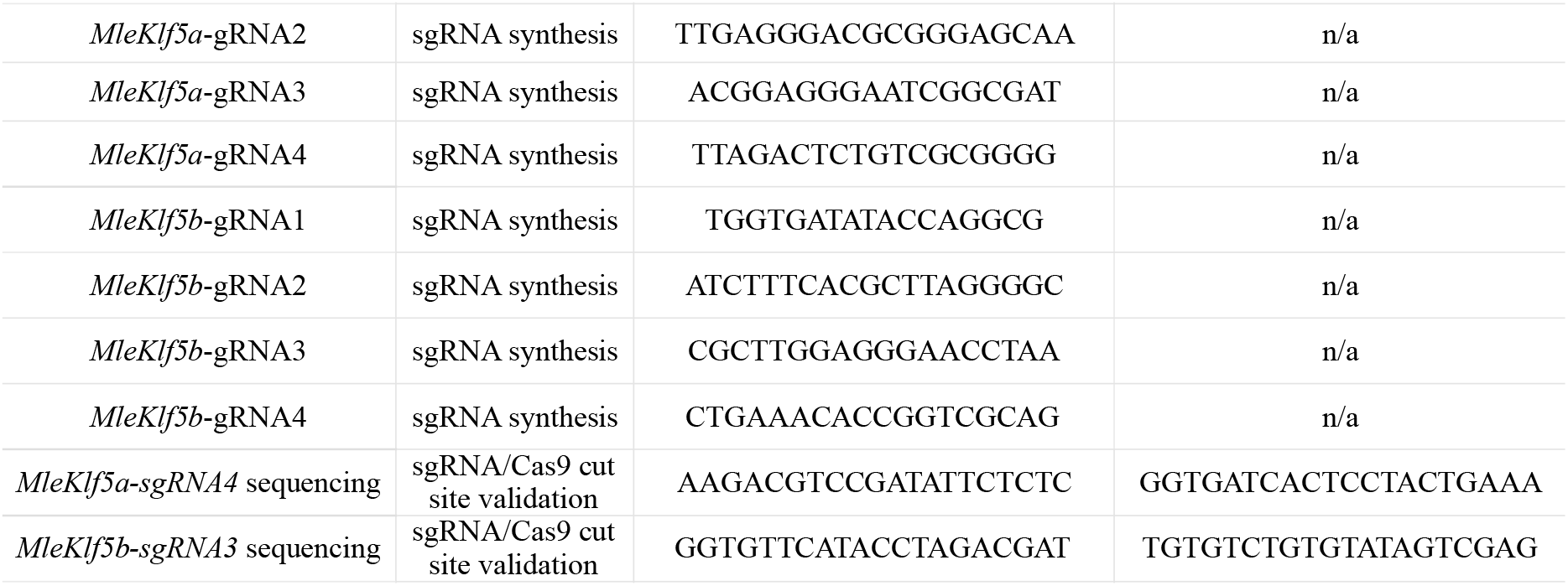
Primers and Oligonucleotides used in this study. All sequences are oriented 5’-3’. Blue nucleotide sequences correspond to T7 promoter. Black bold italicized nucleotide sequences correspond to genomic *MleKlf* targets and include addition of two 5’ *G* residues to aid T7 polymerase binding. Red nucleotide sequences denote region of complementary between templated primers and Universal tracrRNA primer, which are annealed to the form the sgRNA transcription template.

We reduced the chance of potential off target site (OTS) Cas9 mediated exonuclease activity by designing sgRNAs that had no fewer than 3 mismatches to non-target loci in the *M. leidyi* reference genome. To assess potential OTS Cas9 exonuclease activity, we designed primers, amplified and Sanger sequenced regions around the remaining set of predicted low probability cut sites of non-KLF genes. No evidence of Cas9 exonuclease activity was observed (Table 2). Thus, we interpreted that phenotypes generated by both gene abrogation approaches in our study were due to the simultaneous disruption of *MleKlf5a* and *MleKlf5b* gene expression.

**Table 2.**
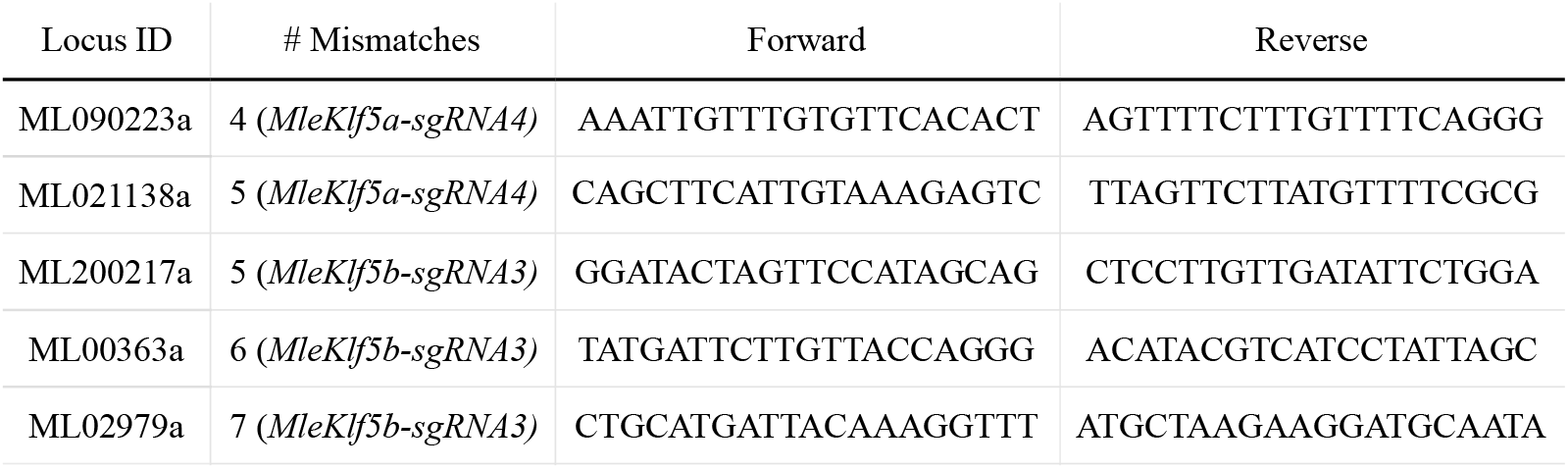
Off-target CRISPR/Cas9 loci with mismatches to either *MleKlf5a* or *MleKl5b* target sequence and primers used for Sanger sequencing. All sequences are oriented 5’-3’.

### Knockdown of zygotic *MleKlf5a* and *MleKlf5b* expression

KLF-MO and KLF-Cas9 embryos phenocopied one another and displayed phenotypes of varying penetrance (Fig. 3A-Q). A higher proportion of severe phenotypes were observed among KLF-Cas9 embryos as compared to KLF-MO embryos (Fig. S2B), reflecting the effects of Cas9-mediated genome editing versus titration of functional mRNAs by sbMOs. In contrast to the observation of predominantly severe phenotypes in KLF-Cas9 embryos injected with *MleKlf5a-sgRNA4*+*MleKlf5b-sgRNA3* (Fig. 3P,Q), single gene knockdown using either *MleKlf5a-sgRNA4* or *MleKlf5b-sgRNA3* primarily generated mild phenotypes (Fig. S3).

**Fig 3.**
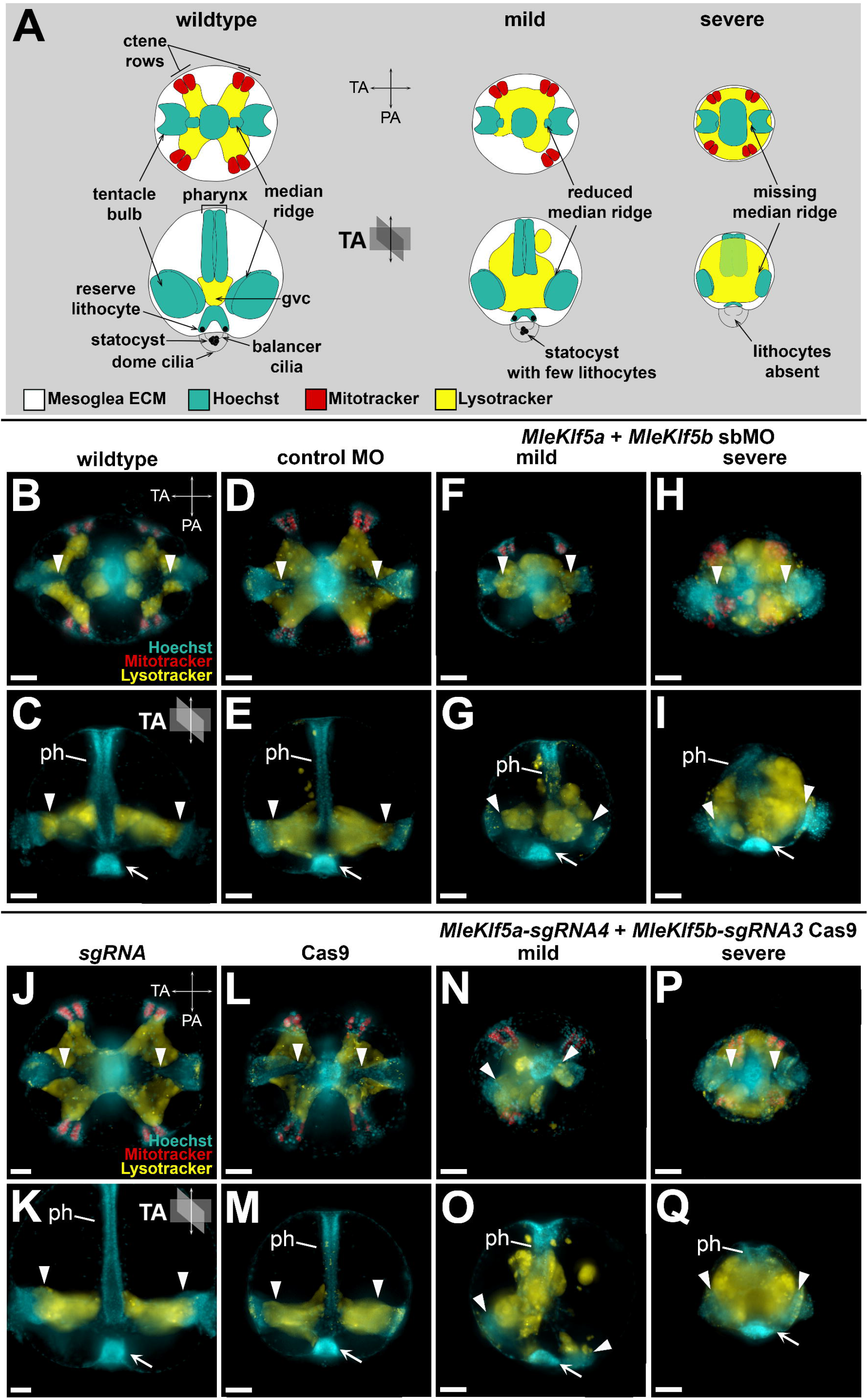
Phenotypes generated by *MleKlf5a* and *MleKlf5b* double gene knockdown via sbMO and *sgRNA*-Cas9 genome editing. (**A)** Representative schematics (see **Fig. 1A**) of wildtype, mild and severe phenotypes highlighting tissues and cell types disrupted in KLF-MO and KLF-Cas9 *M. leidyi* embryos. The top row is an aboral view. The bottom row is a lateral view with oral up and aboral down. (**B-Q**) Representative live images of ∼20 hpf cydippids. Aboral view in **B, D, F, H, J, L, N, P**. Lateral view, oral up, in **C, E, G, I, K, M, O, Q**. Schematic depiction of tentacular axis (TA) and pharyngeal axis (PA) orientation are located in panel upper right. (**B, C**) Un-injected wildtype embryo. Hoechst (blue) marks nuclei. MitoTracker (red) preferentially marks the position of ctene row polster cells, one pair per embryonic quadrant. Lysotracker (yellow) preferentially stains epithelial cells lining the gastrovascular cavity (gvc). Tentacular median ridges (arrowheads) are positioned medially along the tentacular axis and contacted by gvc epithelial cells. The pharynx (ph) is positioned centrally and joins with the gvc aborally. The apical organ (arrow) is located at the aboral pole of the embryo. Morphology is unaffected in embryos sham injected with control morpholino (MO) (**D, E**), *sgRNA* only (**J, K**) or Cas9 protein only (**L, M**). In contrast, mild phenotypes in double gene knockdown KLF-MO embryos (**F, G**) and double gene edited KLF-Cas9 embryos (**N, O**) display aberrant distributions of gvc epithelial cells (Lysotracker signal), aberrant patterning of the pharynx (ph) including aboral bifurcations (**G, O**; refer to Fig. S4), aberrant patterning of the tentacle bulb and tentacular median ridges (arrowheads), and atypical apical organ (arrow) morphology. Severe phenotypes in double gene knockdown KLF-MO embryos (**H, I**) and double gene edited KLF-Cas9 embryos (**P, Q**) are reduced in size due to lack of mesoglea ECM extrusion, display collapsed pharynx with gvc junction defects, significantly reduced tentacle bulbs and tentacular median ridges (arrowheads), and apical organ defects (arrow). Scale bars: 50 µm; aph, ph, pharynx; PA, pharyngeal axis; TA, tentacular axis.

KLF-MO and KLF-Cas9 embryos with mild phenotypes underwent pharyngeal elongation simultaneous with both mesoglea extrusion and a concomitant increase in size similar to that observed in control embryos; however, experimental embryos displayed disorganized patterning at the aboral end of the pharynx and the infundibular gastrovascular cavity (Fig. 3F,G,N,O). Occasionally, we observed pharyngeal bifurcation at the junction of the pharynx with the infundibular gastrovascular cavity (Fig. 3G,O, Fig. S4). In contrast, in severely affected embryos, the internal embryonic space typically occupied by mesogleal extracellular matrix (ECM) was absent and the interior volume was completely occupied by gastrovascular endoderm and abnormally elongated pharyngeal tissue. Thus, embryos having severe phenotypes failed to increase in size most likely due to the lack of ECM extrusion into the mesoglea space (Fig. 3A,H,I,P,Q). Both the stomodeum and oral regions of the pharynx were still visible in severe mutant embryos, indicating that the entire pharyngeal structure was not lost. However, it is unclear whether the observed abnormal pharyngeal elongation is caused by *Klf* gene abrogation directly or is a spatial effect due to the absence of mesogleal ECM.

Among both KLF-MO and KLF-Cas9 embryos, patterning defects were also observed in the apical organ (Fig. 3, Fig. 4A-G). *MleKlf5a* and *MleKlf5b* knockdown resulted in a significant reduction of apical organ lithocytes as compared to control embryos (Fig. 4A-G). By 20 hpf, control embryo statocysts contained an average of ∼7 lithocytes (Fig. 4A,B,G). KLF-MO embryos had an average ∼4 lithocytes, with three embryos lacking lithocytes entirely (Fig. 4C,D,G). KLF-Cas9 embryos had an average of ∼2 lithocytes, with five embryos completely lacking lithocytes (Fig. 4E-G). Notably both KLF-MO and KLF-Cas9 embryos lacking lithocytes still possessed phenotypically normal balancer cilia and dome cilia, tissues derived from ectoderm (Fig. 4D,F).

**Fig 4.**
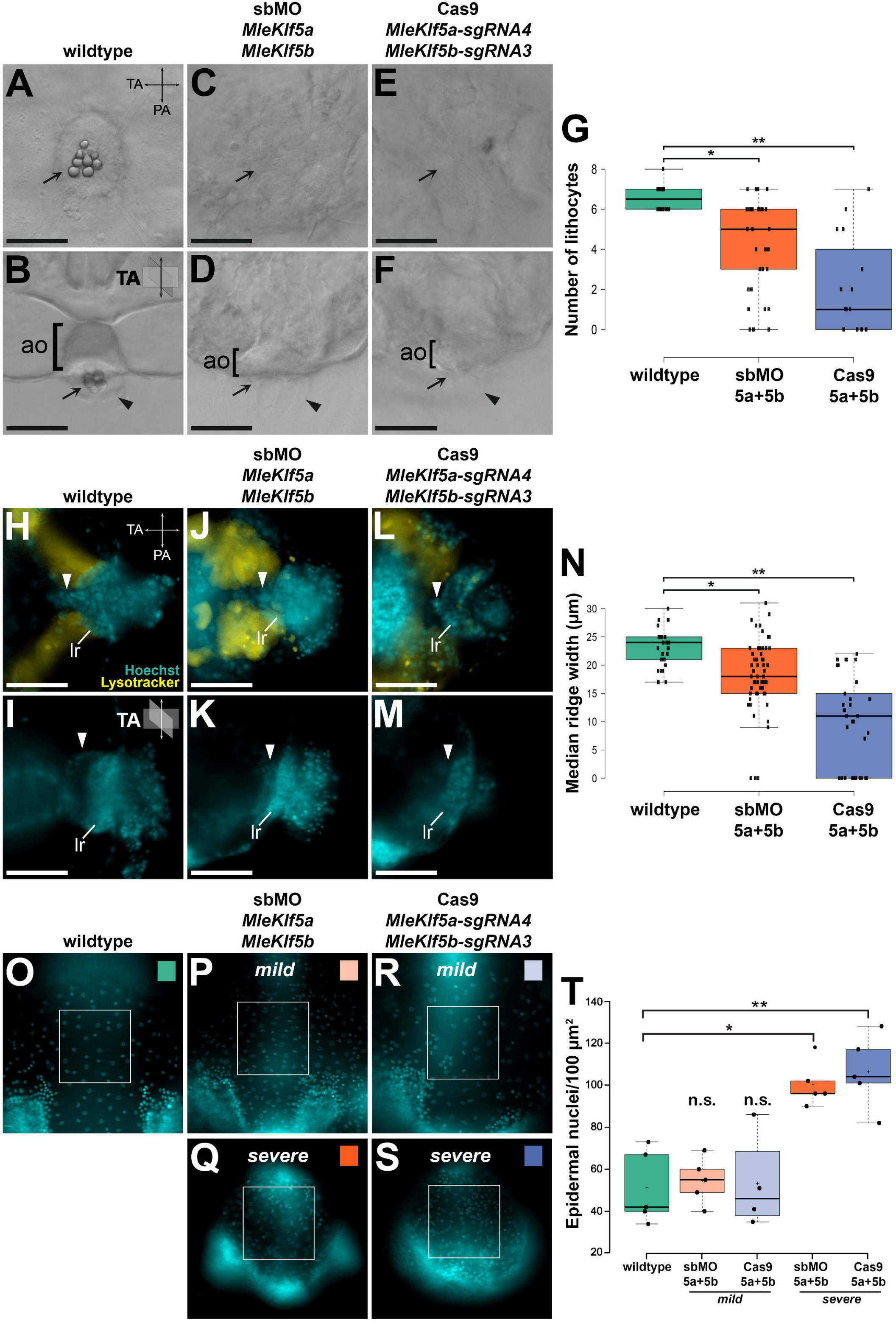
*MleKlf5a* and *MleKlf5b* double gene knockdown disrupts the development of endodermally derived cell types and structures including lithocytes and the tentacular median ridge. Live images of embryos at ∼20 hpf. Schematic depiction of tentacular axis (TA) and pharyngeal axis (PA) orientation are located in panel upper right. Aboral view in **A, C, E, H, J, L**. Lateral view, oral up, in **B, D, F, I, K, M, O-S**. (**A, B**) Wildtype embryo with view of the apical organ (ao) showing position of lithocytes (arrow) and dome cilia (arrowhead). (**C, D**) Representative double gene knockdown KLF-MO embryo and (**E, F**) representative double gene edited KLF-Cas9 embryo lacking lithocytes. Dome cilia (arrowheads) and balancer cilia are present in both KLF-MO and KLF-Cas9 embryos. (**G**) Quantification of lithocyte production. *MleKlf5a* and *MleKlf5b* double gene knockdown significantly reduces lithocyte production. Centerlines show the medians; box limits indicate the 25th and 75th percentiles; whiskers extend 1.5 times the interquartile range from the 25th and 75th percentiles; * = two-tailed *t*-test, *t* = 3.47, *p*<0.005; ** = two-tailed *t*-test, *t* = 6.52, *p*<0.00001. Individual counts are plotted as black dots where n = 14, 33, and 15 embryos, respectively. (**H, I**) Wildtype tentacular median ridge (arrowhead) and lateral ridge (lr). (**J, K**) Representative double gene knockdown KLF-MO embryo and (**L, M**) representative double gene-edited KLF-Cas9 embryo with dramatically reduced tentacular median ridge. The tentacle bulb lateral ridge remains present in both KLF-MO and KLF-Cas9 embryos. (**N**) Quantification of tentacular median ridge width. *MleKlf5a* and *MleKlf5b* double gene knockdown significantly reduces tentacular median ridge width. Centerlines show the medians; box limits indicate the 25th and 75th percentiles; whiskers extend 1.5 times the interquartile range from the 25th and 75th percentiles; * = two-tailed t-test, t = 4.01, p<0.0005; ** = two-tailed t-test, t = 8.32, p<0.00001. Individual measurements are plotted as black dots where n = 28, 61, and 34 tentacular median ridge widths, respectively. Each measurement represents a single tentacular median ridge width, with a maximum of 2 from each embryo (i.e., an individual embryo has two tentacular median ridges, thus each embryo may contribute 2 tentacular median ridge width measurements). A measurement of 0 indicates the absence of a tentacular median ridge and/or tentacle bulb. (**O-S**) Representative images from a subset of each group of embryos with a 100 μm^2^ region of interest focused on the outer epidermal cell layer of wildtype (**O**), KLF-MO mild (**P**) and severe (**Q**), and KLF-Cas9 mild (**R**), and severe (**S**) embryos. (**T**) Quantification of epidermal nuclei cell counts. Centerlines show the medians; box limits indicate the 25th and 75th percentiles; whiskers extend 1.5 times the interquartile range from the 25th and 75th percentiles; crosses represent sample means; **\*** = two-tailed *t*-test, *t*= -5.35225, *p* < .005; **\*\*** = two-tailed *t*-test, *t* = -4.99757, *p* < .005. Area nuclei counts are plotted as black dots where n = 5, 5, 4, 5, and 5 area count samples respectively. KLF-MO severe (*M = 100*.*40, SD = 10*.*7*) and KLF-Cas9 severe (*M = 106*.*40, SD = 17*.*4*) both had a significantly higher density of epidermal nuclei counts per 100 μm^2^ area than control embryos (*M = 51*.*20, SD = 17*.*5*). There was no significant difference in epidermal nuclei counts for KLF-MO mild (*M = 54*.*60, SD = 11*.*0*) or KLF-Cas9 mild (*M = 53*.*25, SD = 22*.*8*) relative to control embryos. Scale bars: 50 µm.

The simultaneous abrogation of *MleKlf5a* and *MleKlf5b* also resulted in a dramatic reduction in tentacle bulb size, particularly in the tentacular median ridge (Fig. 3, Fig. 4H-N). We measured the tentacular median ridge width and found significant differences between control and injected embryos (Fig. 4N, Fig. S5). The control embryo average tentacular median ridge width was ∼23 µm. KLF-MO (Fig. 4J,K) and KLF-Cas9 (Fig. 4L,M) embryo average tentacular median ridge width was ∼18 µm and ∼9 µm, respectively (Fig. 4N). Moreover, we observed that 15% of KLF-MO embryos and 29% of KLF-Cas9 embryos lacked tentacular median ridges altogether (Fig. 4J-N).

In severely affected animals we observed a significant increase in the density of epidermal cells, ∼100 nuclei/100 µm^2^ in severe embryos compared to ∼50 nuclei/100 µm^2^ in wildtype embryos (p < .005; Fig. 4O-T). The spacing between epidermal cell nuclei was closer among the severe phenotypes relative to normally developing animals (Fig. 4O-S). This suggests reduced lateral tension forces on epidermal cells in animals lacking underlying mesogleal ECM and indicates that the total number of epidermal cells remained the same, only their spatial relationship was altered (i.e., closer spacing of nuclei). Thus, despite a decreased total body size, the ectodermal cell contribution to the epidermis appears to be largely unaffected.

The tentacular median ridge in adult *Pleurobrachia pileus* and juvenile *M. leidyi* cydippids has previously been shown to contain populations of proliferative cells (Alié et al., 2011; Reitzel et al., 2016; Schnitzler et al., 2014). In our *MleKlf5a* and *MleKlf5b* knockdown experiments, the relative size of the tentacular median ridge was consistently reduced, therefore we decided to perform EdU incorporation assays during mid-late embryogenesis to assess cell proliferation (Fig. S6). We observed reduced EdU incorporation in areas affected by the knockdown of *MleKlf5a* and *MleKlf5b*, including the tentacular median ridge and pharynx, suggesting that reduced cell proliferation rates are associated with the attenuation of zygotic *Mle*KLF5a and *Mle*KLF5b activity (Fig. S6J,K).

## Discussion

Our expression analyses of *MleKlf5a* and *MleKlf5b* in *M. leidyi* show that transcripts of both genes are maternally loaded and ubiquitously distributed through gastrulation (Fig. 1B-E), corroborating previous RNA-seq results which detected abundant transcripts for both *MleKlf5a* and *MleKlf5b* during *M. leidyi* early embryonic cleavage stages (Davidson et al., 2017). Knockdown of zygotic *MleKlf5a* and *MleKlf5b* expression does not appear to impact early embryonic development, as injected embryos underwent normal early cleavage and gastrulation. The zygotic expression of *MleKlf5a* and *MleKlf5b* display localized spatio-temporal patterns in post-gastrulation embryos and transcripts for both *MleKlf5a* and *MleKlf5b* are expressed in the developing pharynx, gastrodermis, tentacle bulbs and apical organ (Fig. 1F-Q). These similar expression patterns could be due to functionally redundant roles (Lynch and Conery, 2000). In contrast, the expression of *MleKlfX* transcripts are restricted to late stages of development in a subset of apical organ epithelial cells (Fig. S1C,D). The *M. leidyi KlfX* gene sequence is highly divergent relative to other metazoan *Klf* genes (Presnell et al., 2015), suggestive of a *Mnemiopsis-* specific functional role for *MleKlfX*.

The *Klf* gene complement in *M. leidyi* is reduced compared to other non-bilaterian lineages (Presnell et al., 2015), a trend observed in other ctenophore gene families (Moroz et al., 2014; Ryan et al., 2013). *Klf5-like* genes are found in all metazoans (McCulloch and Koenig, 2020; Presnell et al., 2015). Among the non-bilaterian phyla, a *Klf5* ortholog in the cnidarian *Nematostella vectensis* genome was shown to be expressed in a cluster of cells associated with digestive filaments and the gastrodermis (Sebé-Pedrós et al., 2018a). In sponges, a *Klf5* ortholog was found to be expressed in stem-cell like archaeocytes in the marine sponge *Amphimedon queenslandica* (Sebé-Pedrós et al., 2018b), and in the digestive choanocytes and peptidocytes of the freshwater sponge *Spongilla lacustris* (Musser et al., 2019 preprint). In vertebrates, *Klf5* orthologs are required for the maintenance of intestinal crypt epithelia in the gut (Gao et al., 2015; Kuruvilla et al., 2015; McConnell et al., 2011; Nandan et al., 2015). While less is known about *Klf5* orthologs from invertebrate bilaterians, *Klf5* is expressed in several cephalopod embryonic tissues including yolk cells and the developing mouth (McCulloch and Koenig, 2020). In our previous phylogenetic study, it was unclear whether the few identified invertebrate sequences were either *Klf4* or *Klf5*, which share high sequence similarity (Presnell et al., 2015). One of these sequences, *Drosophila melanogaster dar1*, shares sequence similarity to human *Klf5* but has a functional role more similar to human *Klf4*, and was shown to play a role in regulation of gut proliferation (Wu et al., 2018b). Based on our expression analysis of *MleKlf5a* and *MleKlf5b* and the observed dysregulation of gastrodermal patterning in *MleKlf5a+MleKlf5b* knockdown embryos, our data suggest an evolutionarily conserved role for *Klf5-like* orthologs in the regulation and maintenance of gut epithelia among metazoans.

*M. leidyi* endodermal cell lineages stem from early cleavage stage E and M oral macromeres while ectodermal lineages originate from the aboral micromeres. Fate mapping experiments show that the ectodermal micromeres contribute to the epidermis, ctene rows, tentacle epithelia and colloblasts, balancer cilia and the epithelial floor of the apical organ, while the endodermal macromeres give rise to the gastrodermis and associated endodermal canal system, muscle, tentacular median ridge, and apical organ lithocytes (Henry and Martindale, 2001; Martindale and Henry, 1997a; Martindale and Henry, 1999). Dysregulation of *MleKlf5a* and *MleKlf5b* show consistent abnormal phenotypes associated with the development of the apical organ and tentacle bulbs. In the apical organ of *MleKlf5a* and *MleKlf5b* dysregulated embryos, the development of endodermally derived lithocytes is reduced or absent while the ectodermally derived epithelial floor, balancer cilia, and dome cilia appear normal (Fig. 4A-G). Similarly in the developing tentacle bulb, abrogation of *MleKlf5a* and *MleKlf5b* activity resulted in the absence or reduction in size of the endodermally derived tentacular median ridge, which gives rise to the tentacle muscular core (Alié et al., 2011; Fig. 4H-N, Fig. S6L-N). Remaining tentacle tissue likely represents ectodermal contributions to tentacle epithelia and colloblasts. The development of other ectodermally derived structures, including the stomodeum and epidermal cells (Martindale and Henry, 1999), were unaffected (Fig. 4O-T). These results suggest that *MleKlf5a* and *MleKlf5b* play a functional role in the development and maintenance of endodermally derived tissues during *M. leidyi* embryogenesis.

With regard to the unique ectodermal expression domain of *MleKlf5b* (Fig. S1A,B), overall no ectodermal or ctene row patterning phenotypes were observed in KLF-Cas9 embryos. In a few cases, ctene rows showed gross spatial disorganization, possibly reflecting a requirement for coordinated contact between ectoderm and underlying endoderm for precise ctene row alignment. For example, in phenotypically mild KLF-MO and KLF-Cas9 embryos, ctene row morphogenesis did not occur in quadrants in which endodermal tissue failed to contact ectodermal tissue (Fig. 3A,F,N). This result corroborates prior analyses indicating that ctene row development is at least partially regulated through inductive interactions between endodermal and ectodermal cell lineages (Fischer et al., 2014; Henry and Martindale, 2001; Henry and Martindale, 2004; Martindale and Henry, 1997a). One possible explanation for the observed *MleKlf5b* expression pattern could be that *MleKlf5b* is expressed in developing light producing photocytes derived from endodermal 2M macromeres that run subjacent to the ctene rows (Anctil, 1985; Fischer et al., 2014; Freeman and Reynolds, 1973; Martindale and Henry, 1999; Schnitzler et al., 2012). An EdU-positive ring of proliferative cells is situated around the ctene rows (Fig. S6C,G). These proliferative, *MleKlf5b* positive cells may represent photocyte progenitor cells, as photocytes differentiate relatively early during development (Fischer et al., 2014). Notably, the initial development of differentiated polster cells/ctenes is specified by maternal factors, with additional ctenes generated post embryonically. Therefore, zygotic *MleKlf5b* would not directly impact the specification of the initial ctenes during the stages observed in our study. An alternative explanation is that these *MleKlf5b*- and EdU-positive ectodermal cells represent progenitor cells that will give rise to new polster cells post-hatching and thus contribute to ctene row expansion.

In mammalian lineages *Klf5* orthologs help maintain stem cell renewal and promote proliferation in the intestinal crypt and in pluripotent embryonic stem cells (Jiang et al., 2008; Kuruvilla et al., 2015; Nandan et al., 2015; Parisi et al., 2008; Parisi et al., 2010). However, a recent study suggests that mammalian pluripotency factors are not necessarily conserved in all animals, and the ancestral metazoan stem cell toolkit primarily consisted of genes associated with the germline multipotency program (Alié et al., 2015; Juliano et al., 2010). Germline genes, including *Piwi, Bruno*, and *Pl-10*, have been shown to be expressed in putative progenitor cell populations in the tentacle bulb, ctene rows, and apical organ of adult *Pleurobrachia* (Alié et al., 2011). In *M. leidyi* cydippids, *Piwi, Vasa*, as well as *Sox* pluripotency factors are expressed in these same tissues, suggesting that progenitor cell populations in these tissues express both pluripotency factors as well as germline factors (Reitzel et al., 2016; Schnitzler et al., 2014). Our EdU-staining recapitulates earlier work identifying zones of cell proliferation associated with the developing pharynx, gastrodermis, areas around the ctene rows, and in the apical organ epithelial floor (Reitzel et al., 2016; Schnitzler et al., 2014; Fig. S6B-I). These areas of cell proliferation correlate with the zygotic transcript expression domains, including the tentacular median ridge, of both *MleKlf5a* and *MleKlf5b* (Fig. 1, Fig. S6J).

Notably, sponge orthologs to *Klf5, Piwi, Bruno* and *Pl-10* are expressed in archaeocyte and choanocyte cell types variably recognized as sponge equivalents to totipotent, pluripotent, and/or multipotent stem cells (Alié et al., 2015; Musser et al., 2019 preprint; Nakanishi et al., 2014; Sebé-Pedrós et al., 2018b; Sogabe et al., 2019). Although we were unable to perform quantitative analyses, our qualitative assessments show a diminution/loss of EdU-positive cells in the tentacular median ridge and apical organ in *MleKlf5a*+*MleKlf5b* knockdown embryos (Fig. 6K). One interpretation of our results is that *MleKlf5a* and *MleKlf5b* are expressed in proliferative cells and play a functional role in the maintenance of multipotent endodermal progenitor cell populations.

To resolve whether *MleKlf5a* or *MleKlf5b* expressing cells are both proliferative and multipotent will require additional experimentation. Future experiments involving the knockdown of pluripotency and germline determination genes, such as *Piwi* and *Vasa*, along with EdU assays may reveal further aspects of cellular proliferation and specification associated with *Klf* activity. Alternatively, the observed phenotypes may be due to proliferation-independent mechanisms establishing terminal cell identity. For example, *MleKlf5a* and *MleKlf5b* may regulate the terminal specification of lithocyte and tentacle muscle cell types. Based on this work, while the explicit regulatory role of *MleKlf5a* and *MleKlf5b* remains unclear, our results show that *MleKlf5a* and *MleKlf5b* are functionally associated with the formation, developmental patterning and maintenance of endodermally derived structures in *M. leidyi* including the gastrodermis, the tentacular median ridge, tentacle muscle, and apical organ lithocytes. This functional activity may be through the maintenance of multipotent progenitor cell proliferation, and may represent a conserved ancestral function for this transcription factor gene family in the animal stem lineage. Overall, our results begin to lay the groundwork for assessing gene function essential for the embryonic development of *M. leidyi* and thus inform developmental mechanisms unique to Ctenophora for the specification of terminally differentiated tissue and cell types (e.g., lithocytes).

In this study, we report the first use of CRISPR/Cas9 mutagenesis to investigate gene function in a species of ctenophore. We describe techniques that are cost effective and can easily be used by others to assess phenotypes and validate Cas9 activity (e.g., vital dye labeling, Sanger sequencing). This foundational work shows that CRISPR/Cas9 is an effective method for evaluating developmental phenotypes from single or combinatorial gene function loss in G0 ctenophore embryos. Future studies can refine our protocol to generate more efficient CRISPR/Cas9 mutagenesis by choosing different targets within loci (e.g., the transcriptional start site) and by increasing or modifying Cas9 exonuclease activity. Techniques have recently been developed that improve Cas9 editing, resulting in high percentages (>80%) of indel mutations (Hoshijima et al., 2019; Wu et al., 2018a). Although cell-autonomous phenotypes can be detected in G0 Cas9-injected embryos, which is useful for generating hypotheses regarding gene function, the characterization of stable and heritable non-lethal mutations (i.e., in F1 embryos) would be even better. *M. leidyi* are self-fertile hermaphrodites which could be leveraged to enable rapid creation of stable lines useful for characterization of mutations generated via CRISPR/Cas9. Along with recent RNA-seq data highlighting candidate genes associated with zygotic gene activation and patterning of specific cell types in ctenophores (Babonis et al., 2018; Davidson et al., 2017; Sebé-Pedrós et al., 2018b), CRISPR/Cas9 mutagenesis in *M. leidyi* (and potentially other ctenophore species) will provide much needed insight into the genetic mechanisms underlying unique facets of ctenophore biology (Bessho-Uehara et al., 2020; Jokura et al., 2019; Yamada et al., 2010) and further our understanding of early metazoan evolution.

## Materials and methods

### Cloning and *in situ* hybridization

RNA was extracted using Trizol (Thermo Fisher Scientific) from *Mnemiopsis* embryos collected at different developmental stages and used to generate cDNA libraries (SMARTer kit, Clontech). The coding sequences of *MleKlf5a, MleKlf5b*, and *MleKlfX* were amplified from cDNA (Table 1) and cloned into pGEM-T Easy vector (Promega). The cloned fragments were used as templates for *in vitro* transcription (MEGAscript, Ambion) of antisense digoxigenin-labeled (Digoxigenin-11-UTP, Roche) riboprobes.

*In situ* hybridization followed (Pang and Martindale, 2008). Riboprobes were used at a final concentration of ∼0.5 ng/µl and hybridized with embryos for 24 hours. After color development, nuclei were labeled with either DAPI (Molecular Probes) or Hoechst 33342 (Molecular Probes) in 1x PBS. Embryos were immediately imaged or stored at -20°C in 70% glycerol in 1x PBS. Images were acquired using a Zeiss Axio Imager.Z2, Zeiss AxioCam MRm Rev3 camera, and Zeiss Zen Blue software. Fluorescent Z-stacks were deconvolved, post-processed for brightness and contrast and assembled in Adobe Photoshop. Monochrome brightfield images were inverted, pseudo colored and overlaid onto fluorescent images of labeled nuclei.

### EdU labeling

Click-iT® EdU Alexa Fluor® 647 Imaging Kit (ThermoFisher Scientific) was used for identification of proliferating cells. Embryos were collected at different developmental stages and pulse incubated for 25 minutes with 100 µM EdU in a solution of a 1:1 volumetric ratio of artificial seawater (FSW) to 6.5% MgCl2 (dissolved in dH2O) at room temperature. The EdU solution was washed out and embryos were either fixed immediately or allowed to continue to develop during a 24-hour chase and subsequently fixed. Embryos were fixed with 4% PFA in FSW for 30 minutes at room temperature, washed with 3% BSA in 1x PBS, and incubated with 0.5% Triton X-100 in 1x PBS for 20 minutes at room temperature. Fixed embryos were washed with 3% BSA in 1x PBS and stored at 4°C until used for EdU detection as per manufacturer protocol. Embryos were subsequently washed with 1x PBS and mounted on glass microscope slides. Images were acquired using a Zeiss Axio Imager.Z2, Zeiss AxioCam MRm Rev3 camera, and Zeiss Zen Blue software. Fluorescent Z-stacks were deconvolved, post-processed for brightness and contrast, and assembled in Adobe Photoshop or FIJI (Schindelin et al., 2012).

### Preparation and microinjection of embryos

Microinjection needles were pulled with a Brown micropipette puller (P-1000, Sutter Instrument Company) using filamented aluminosilicate glass capillaries (AF100-64-10, Sutter Instrument Company). Pulled capillary needles were beveled using a microelectrode beveler (BV-10, Sutter Instrument Company). Beveling creates a consistent microinjection needle with uniform tip characteristics optimized for egg penetration and substantially reduces embryo mortality. Beveled capillary needles were loaded via backfilling with injection cocktails mixed with fluorescently-conjugated dextran (Invitrogen) for rapid assessment of injection success and subsequent lineage tracing. Loaded capillary needles were mounted to a Xenoworks microinjection system (Sutter Instrument Company) paired to a Zeiss Discovery V8 epifluorescence stereomicroscope.

Microinjection dishes were designed to aid in stabilizing and positioning embryos during injections. In a 30 mm or 60 mm petri dish, a glass microscope slide was placed at a 30-45° angle. Molten 2% agarose (dissolved in 1:1 volume FSW:dH2O) was slowly poured into the dish until the agarose meniscus reached the underside of the angled glass slide. Once the agarose solidified, the glass slide was removed, creating a molded ramp impression terminating in a 90° trough. For short-term storage of agarose molds between microinjection sessions, we flooded dishes with 1x Penicillin/Streptomycin:FSW (PS:FSW), sealed and stored at 4°C.

Laboratory cultures of adult *Mnemiopsis leidyi* on a ∼12 hr:12 hr light:dark cycle were spawned ∼4 hours post darkness (hpd). At ∼3.5 hpd individual adult *M. leidyi* were placed into 8-inch glass bowls (Carolina Biological Supply) and screened for mature sperm and eggs. Freshly fertilized eggs were collected by pipette and passed sequentially through a 500 µm and a 400 µm cell strainer (pluriSelect Life Science) to remove excess mucus and egg jelly. Embryos were then washed with PS:FSW. Ctenophore vitelline membranes are resistant to penetration from microinjection needles and must be removed. Additionally, the highly viscous inner egg jelly, which will clog the injection needle, should be removed from the egg surface. In gelatin-coated dishes filled with PS:FSW, we used acid sharpened tungsten needles to remove both the vitelline membranes and underlying egg jelly. A 5x gelatin stock (0.5% Knox unflavored gelatin dissolved in dH2O, then formalin added to a final concentration of 0.19%) was diluted to 1x with dH2O, poured into dishes and swirled, and then discarded. Once the gelatin dried, the dishes were rinsed several times with dH2O. Applying a gelatin coating helps prevent devitellinized embryos from adhering to plastic, glass and metal surfaces. We applied a gelatin coat to glass and plastic dishes, transfer pipettes, and dissecting needles. Once the vitelline membranes and egg jelly were removed, embryos were then carefully transferred to an injection dish and positioned along the agarose trough for microinjection. After injections, embryos were kept at room temperature in gelatin-coated dishes until reaching the desired development stage for further analyses.

### Morpholino oligonucleotides

Splice-blocking morpholino oligonucleotides (sbMOs, Gene Tools) were designed for both *MleKlf5a* (*ML00922a*) and *MleKlf5b* (*ML25776a*). *MleKlf5a* sbMO #1 and sbMO #2 targeted intron 2-exon 3 and intron 3-exon 4 boundaries, respectively. *MleKlf5b* sbMO #1 and sbMO #2 targeted exon 6-intron 6 and exon 7-intron 7 boundaries, respectively. A standard control MO was used as a negative control. Sequences of sbMOs are listed in Table 1. Stock solutions of sbMO in dH2O were stored at room temperature. sbMO injection cocktail solutions consisted of a final sbMO concentration of ∼333 nM and ∼0.5 mg/ml fluorescent dextran (rhodamine or Alexa-Fluor 488, 10,000 MW, Invitrogen) in 35% glycerol. After phenotypic analyses via vital-dye staining and microscopy, RNA was extracted from individual embryos (Arcturus PicoPure, ThermoFisher) and cDNA prepared. Gene-specific primers were used on cDNA (OneTaq One-Step RT-PCR, New England Biolabs) to evaluate aberrant transcript splicing via gel electrophoresis. A total of 45 embryos were injected with a *MleKlf5a* + *MleKlf5b* double-gene knockdown sbMO cocktail and used for all downstream analyses.

### CRISPR/Cas9 mutagenesis

We followed a cloning-free method to generate sgRNAs (Kistler et al., 2015; Varshney et al., 2015). PCR amplified templates were generated by annealing a 20-nt universal tracrRNA oligo to a sgRNA-specific oligo that consisted of a T7 promoter, followed by the sgRNA target sequence, and a complementary sequence to the tracrRNA oligo (Table 1). These templates were then *in vitro* transcribed (MEGAscript, Ambion) to generate sgRNAs. The CasOT program (Xiao et al., 2014) and *M. leidyi* reference genome (Moreland et al., 2014; Moreland et al., 2020) were used to identify sgRNA target sites for *MleKlf5a* (*ML00922a*), *MleKlf5b* (*ML25776a*), and *MleKlfX* (*ML20061a*). We selected sgRNAs that had no fewer than four mismatches to alternative genomic sites to minimize potential off-target site (OTS) activity (Table 1; Table 2). Recombinant Cas9 protein (PNA Bio) and sgRNAs were injected at concentrations of 400 ng/µl of Cas9 protein and 100 ng/µl for each sgRNA. A total of 17 embryos from *MleKlf5a* + *MleKlf5b* double-gene knockout sgRNA/Cas9 cocktail injections were live imaged and processed for downstream analyses. After phenotypic analysis, genomic DNA was extracted from individual embryos (QIAamp DNA Micro, Qiagen) and each sgRNA target site was amplified and Sanger sequenced. The ICE analysis tool (Hsiau et al., 2019 preprint) was used to determine Cas9 efficiency for each sgRNA. ICE analysis gives two scores: an ICE score which reflects the percentage of indels found and a KO score which reflects the percentage of indels that produce a frameshift mutation. We obtained ICE/sequencing information and analyzed off target sites from all 17 embryos for *MleKlf5b* cut sites and 14 of the 17 embryos for *MleKlf5a* cut sites. Additional single-gene injection analyses, either *MleKlf5a* or *MleKl5b*, were performed on five embryos per gene.

### Phenotypic analysis through vital dye staining

Control, sbMO, and Cas9 injected embryos at 20-24 hpf were incubated in filtered seawater (FSW) containing a final concentration of 100 nM MitoTracker (Deep Red FM, Molecular Probes), 100 nM LysoTracker (Red DND-99, Molecular Probes), and 10 ng/µl Hoechst 33342 for one hour at room temperature. The live embryos were then placed on glass slides in a drop of FSW and relaxed with a drop of 6.5% MgCl2 (in dH2O) on a coverslip positioned with clay feet for imaging. DIC and fluorescent images were acquired using a Zeiss Axio Imager.Z2, Zeiss AxioCam MRm Rev3 camera, and Zeiss Zen Blue software. Fluorescent Z-stacks were deconvolved, post-processed for brightness and contrast, and assembled in Adobe Photoshop.

### Epidermal nuclei counts

A subset of live images from wildtype, KLF-MO, and KLF-Cas9 embryos were used (see previous section) to quantitate epidermal nuclei. Individual Z-sections from Hoechst channels were focused on the outer epidermal layer for each embryo oriented along the tentacular axis (TA). A 100 μm^2^ region of interest (roi) was positioned medially and oral of the ctene rows. Nuclei within the roi were manually counted. Nuclei counts were quantified and plotted using R (http://shiny.chemgrid.org/boxplotr/).

## ACKNOWLEDGEMENTS

This work was supported in part by startup funds from the University of Miami College of Arts and Sciences to WEB. JSP was supported by the University of Miami College of Arts and Sciences. We thank Ricardo Cepeda for additional animal support and anonymous reviewers for their time and generous feedback.

## AUTHOR CONTRIBUTIONS

WEB originally conceived the study and designed the research. JSP and WEB performed experiments, collected and analyzed data, and wrote the manuscript. JSP and WEB read and approved the final manuscript.

## AUTHOR INFORMATION

Correspondence and requests for materials should be addressed to WEB.

