## Supplemental Figures for "*Krüppel-like factor* gene function in the ctenophore *Mnemiopsis leidyi* assessed by CRISPR/Cas9-mediated genome editing"

Fig. S1

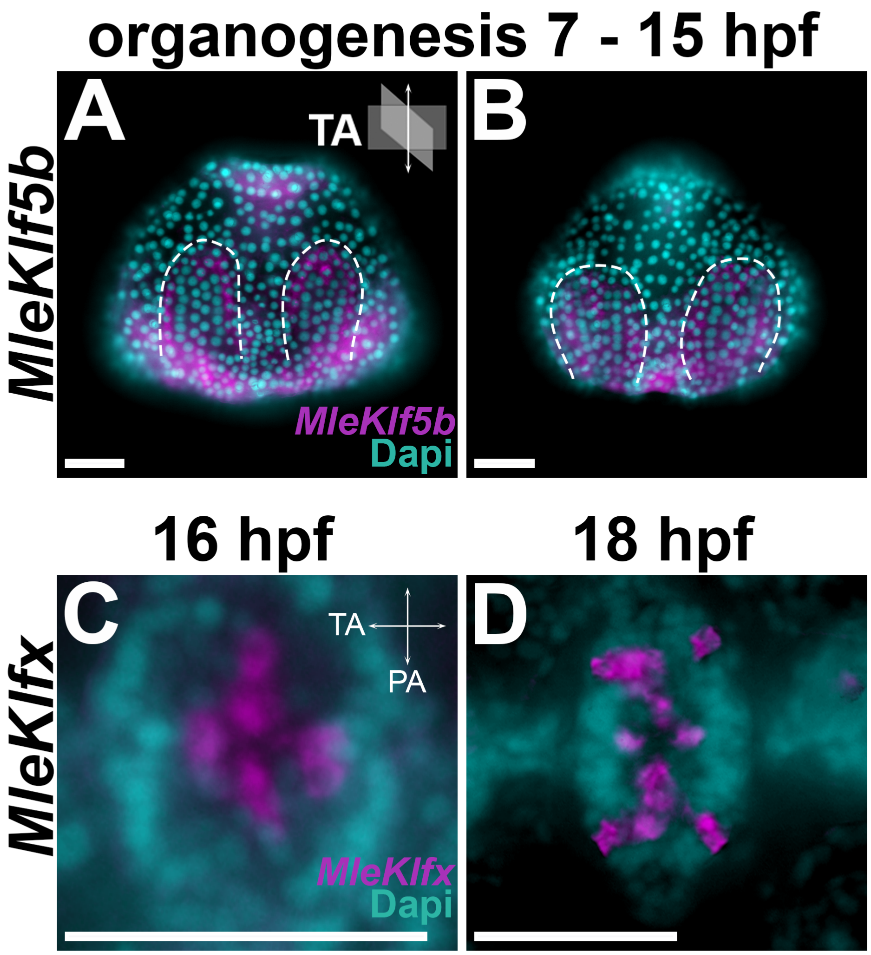


**Figure S1. Additional *MleKlf5b* and *MleKlfX* transcript expression domains.** Schematic depiction of tentacular axis (TA) and pharyngeal axis (PA) orientation are located in panel upper right. (**A, B**) Organogenesis stage 7-15 hpf embryos, lateral view, oral up. Dashed lines bound *MleKlf5b* ectodermal expression (magenta) in rapidly dividing cells (compare with **Fig. S6C, G**) that flank developing ctene row pairs (visible as aligned columns of DAPI stained nuclei bounded by *MleKlf5b* expression). (**C, D**) Magnified aboral views of *MleKlfX* expression (magenta) in epithelial floor cells of the apical organ. (**C**) Initially *MleKlfX* expression is detected at 16 hpf in four small clusters of cells at the boundary of each developing embryonic quadrant converging in the center of the apical organ. (**D**) Expression of *MleKlfX* resolves into several cell clusters by 18 hpf in both the tentacular (TA) and pharyngeal (PA) axes. Scale bars: 50 µm.

**Fig. S2**

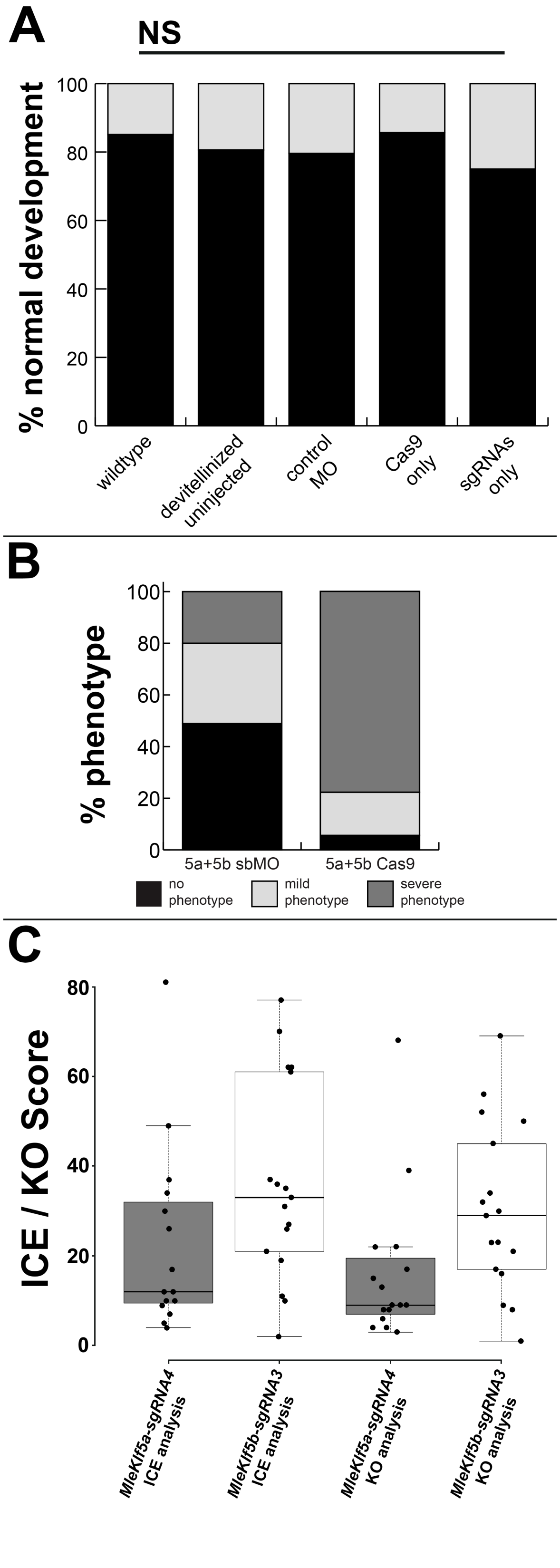


**Figure S2. Injection quantification and ICE analysis of KLF-Cas9 embryos.** (**A**) Sham injection has no effect on embryonic development. Bar graph comparison of percentage of normal development in wildtype $n$ = 161 (24 abnormal); de-vitellinized but uninjected $n$ = 217 (42 abnormal), two-tailed Fisher's exact test $P$=0.2761; control MO injected $n$ = 49 (10 abnormal), two-tailed Fisher's exact test $P$=0.379; Cas9 only injected $n$ = 7 (1 abnormal), two-tailed Fisher's exact test $P$=1; sgRNA only injected $n$ = 4 (1 abnormal), two-tailed Fisher's exact test $P$=0.4851. **(B)** Comparison of phenotypic proportions in KLF-MO embryos (51% no phenotype, 29% mild phenotype, 20% severe phenotype, $n$ = 45) vs KLF-Cas9 embryos (6% no phenotype, 19% mild phenotype, 75% severe phenotype, $n$ = 17). (**C**) Distribution of ICE scores (estimate of indel proportion in signal trace from genomic DNA of individual KLF-Cas9 embryos) and KO scores (estimate of indel proportion that result in frameshift mutation) for *MleKlf5a-sgRNA4* ($n$ = 14) and *MleKlf5b-sgRNA3* ($n$ = 17). Cas9 cut sites were PCR amplified from genomic DNA prepared from individual KLF-Cas9 embryos and subsequently Sanger sequenced. Each cut site data point represents Sanger trace analyses from an individual embryo. Box plot center lines show the median, box limit at 25th and 75th percentiles, whiskers extend 1.5 times interquartile range from the 25th and 75th percentiles, data points are plotted as open circles.

**Fig. S3**

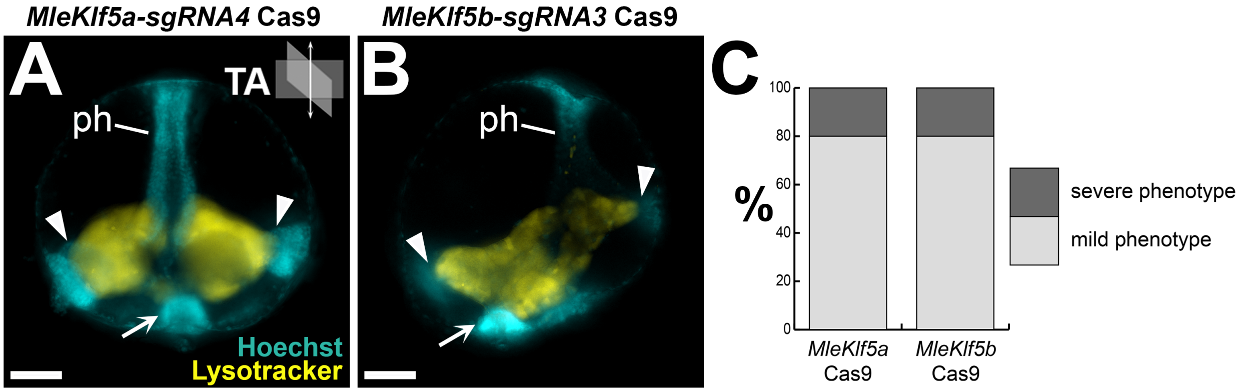


**Figure S3. Cas9 mediated genome editing of either *MleKlf5a* or *MleKlf5b* resulted in reduced phenotype penetrance.** Schematic depiction of tentacular axis (TA) orientation located in panel upper right. Lateral view, oral up. Single gene editing produced primarily mild phenotypes for both *MleKlf5a* KLF-Cas9 embryos (**A**) and *MleKlf5b* KLF-Cas9 embryos (**B**). (**C**) Bar graph comparison of distribution of mild (gray) vs severe (black) phenotypes in *MleKlf5a* KLF-Cas9 embryos ($n$ = 5, 80% mild) and *MleKlf5b* KLF-Cas9 embryos ($n$ = 5, 80% mild). Scale bars: 50 µm.

Fig. S4

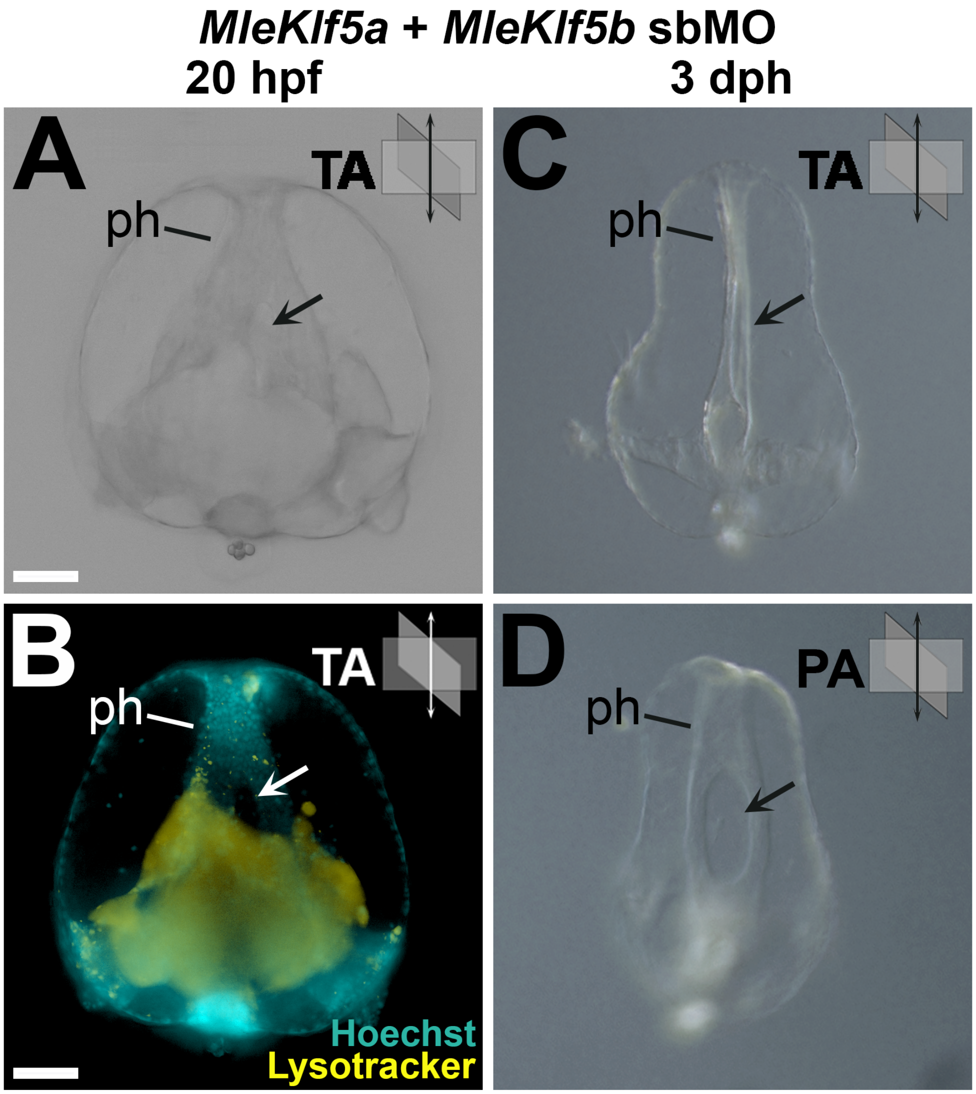


**Figure S4. Aberrant pharyngeal patterning.** Schematic depiction of tentacular axis (TA) and pharyngeal axis (PA) orientation located in panel upper right. Lateral views, oral up. (**A, B**) 20 hpf double gene KLF-MO embryo with bifurcated (arrow) pharynx (ph). (**C, D**) Same individual KLF-MO embryo 3 days post fertilization (dph) showing persistence of pharyngeal bifurcation patterning defect just anterior of the pharyngeal-gastrovascular junction resulting in a deletion of the pharyngeal folds. Scale bars: 50 µm.

**Fig. S5**

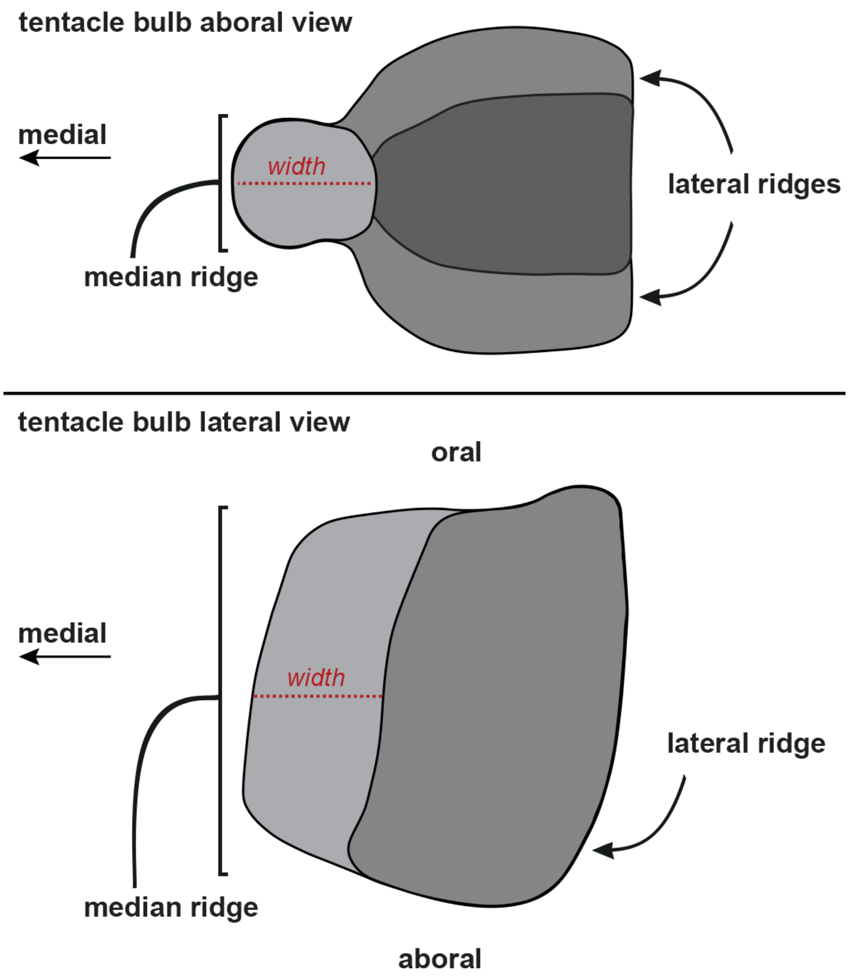


**Figure S5. Cydippid tentacle bulb schematic.** Aboral view: top; Lateral view: bottom. Tentacular median ridge colored light gray. Tentacular lateral ridges colored darker gray. Tentacle (not shown) is rooted at the aboral medial base of the tentacle bulb. Location of measurement taken for the tentacular median ridge width (refer to **Fig. 4N**) is denoted with a red dashed line.

### Fig. S6

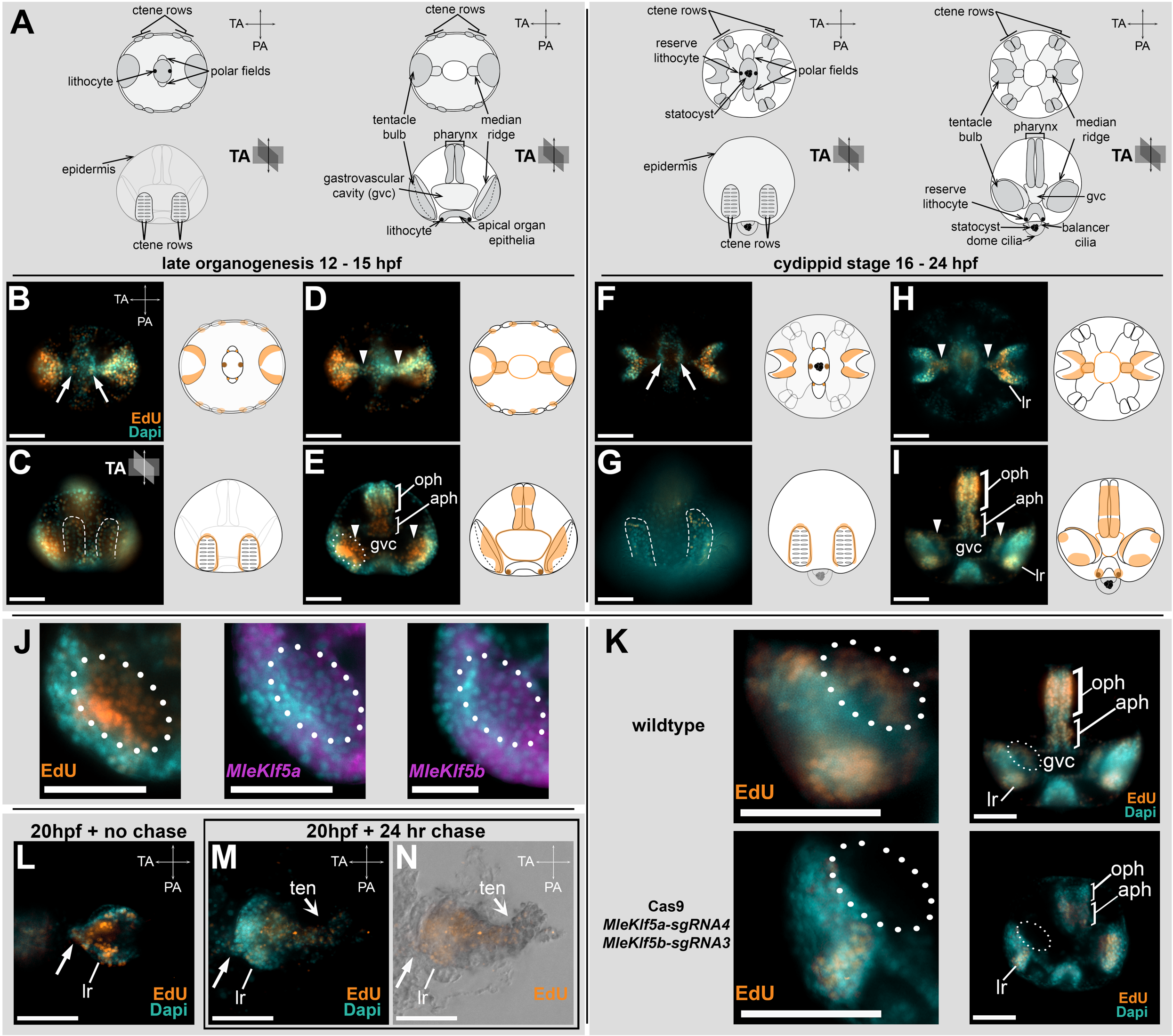


**Figure S6. Regions of rapid cell proliferation.** (**A**) Schematics highlighting major morphological landmarks (e.g., ctene rows, pharynx, tentacle bulbs, apical organ) during *M. leidyi* embryogenesis. The top row is an aboral view. The bottom row is a lateral view with oral up and aboral down. Schematic depiction of tentacular axis (TA) and pharyngeal axis (PA) orientation are located in panel upper right. (**B-I**) Edu incorporation after 25 min pulse during embryogenesis. Orientation follows schematics from **A**. Aboral views in **B,D,F,H,L**. Lateral views in **C,E,G,I,J,K**. EdU incorporation is localized to two cell clusters in the apical organ (**B,F**), epithelial cells flanking the developing ctene rows bounded by dashed lines (**C,G**), epithelial cells lining the gastrovascular cavity (gvc) (**E,I**) in the developing tentacular median ridges (arrowheads), and the oral (oph) and aboral (aph) regions of the pharynx (**E**). (**H,I**) Later in development, EdU incorporation is found in the tentacular median ridge (arrowheads) and lateral ridges (lr) of the tentacle bulbs and in both oral (oph) and aboral (aph) regions of the pharynx. (**J**) Region of interest centered on developing tentacle bulbs from ~14 hpf embryos, showing EdU labeling and *MleKlf5a* and *MleKlf5b* gene expression within the presumptive tentacular median ridge (outlined). (**K, left column**) Region of interest centered on ~20 hpf wildtype and KLF-Cas9 embryo tentacle bulbs highlighting loss of EdU labeling in the tentacular median ridge (outlined) in mutants. **(K, right column)** Whole mount ~20 hpf wildtype and KLF-Cas9 embryo ($n = 2$) showing loss of EdU signal in aboral and oral portions of the pharynx (aph, oph), apical organ, and tentacular median ridge of tentacle bulbs (dashed outline). Note that lateral ridge (lr) cell proliferation is mostly unaffected in KLF-Cas9 embryos. (**L**) Tentacle bulb after 25 min pulse:0 min chase. (**M,N**) Tentacle bulb after 25 min pulse:24 hr chase - EdU incorporation is primarily detected in emergent tentacle muscle cells. Scale bars: 50 µm. TA, tentacular axis; PA, pharyngeal axis.
